# Altered hippocampal place cell representation and theta rhythmicity following moderate prenatal alcohol exposure

**DOI:** 10.1101/2020.02.11.944173

**Authors:** Ryan E. Harvey, Laura E. Berkowitz, Daniel D. Savage, Derek A. Hamilton, Benjamin J. Clark

## Abstract

Prenatal alcohol exposure (PAE) leads to profound deficits in spatial memory and synaptic and cellular alterations to the hippocampus that last into adulthood. Neurons in the hippocampus, called place cells, discharge as an animal enters specific places in an environment, establish distinct ensemble codes for familiar and novel places, and are modulated by local theta rhythms. Spatial memory is thought to critically depend on the integrity of hippocampal place cell firing. We therefore tested the hypothesis that hippocampal place cell firing is impaired after PAE by performing in-vivo recordings from the hippocampi (CA1 and CA3) of moderate PAE and control adult rats. Our results show that hippocampal CA3 neurons from PAE rats have reduced spatial tuning. Secondly, CA1 and CA3 neurons from PAE rats are less likely to orthogonalize their firing between directions of travel on a linear track and between contexts in an open arena compared to control neurons. Lastly, reductions in the number of hippocampal place cells exhibiting significant theta rhythmicity and phase precession were observed which may suggest changes to hippocampal microcircuit function. Together, the reduced spatial tuning and sensitivity to context provides a neural systems-level mechanism to explain spatial memory impairment after moderate PAE.

## Introduction

Prenatal alcohol exposure (PAE) is detrimental to the developing nervous system (Guerri and Sanchis, 1985) and remains one of the most common developmental insults (Day et al., 2002; Green et al., 2009; Thomas et al., 1998). Globally, 9.8% of individuals consume alcohol while pregnant (Popova et al., 2017). There is a large body of evidence supporting the conclusion that even a moderate amount alcohol exposure during fetal development (blood alcohol concentration (BAC) 7-120mg/dl) can disrupt the synaptic and cellular networks believed to underlie episodic and spatial memory resulting in significant lifelong deficits in the ability to remember locations and effectively navigate (Berman and Hannigan, 2000; Harvey et al., 2019; Mira et al., 2020; Sutherland et al., 2000; Valenzuela et al., 2012).

The hippocampus is critical for the encoding of spatial (Morris et al., 1982) and episodic memories (Eichenbaum et al., 2007; Scoville and Milner, 1957). The hippocampal (HPC) subregions dentate gyrus, CA1, CA2, and CA3 contain neurons called “place cells” that selectively fire at particular positions of an animal’s path resulting in receptive fields for particular locations in space (O’Keefe, 1976; O’Keefe and Dostrovsky, 1971). These receptive fields are referred to as place fields and their precise population activity has been thought to form a “cognitive map” giving an animal a sense of past, present, and future locations (O’Keefe and Nadel, 1978). While the correlation between a place cell’s firing rate and location is commonly referred to as rate coding, phase or temporal coding co-occurs whereby spikes from single theta-rhythmic place cells systematically phase shift across successive theta cycles in a phenomenon known as theta phase precession (O’Keefe and Recce, 1993).

Moderate PAE during the first and second trimester impairs spatial behavior (reviewed in Harvey et al., 2019). In the Morris water task, rats with moderate PAE are generally unable to quickly adapt to changes in task demands, such as a change in the escape location in the water task or recalling a previously learned location (Hamilton et al., 2014; Rodriguez et al., 2016; Savage et al., 2002; Sutherland et al., 2000). In dry land tasks, moderate PAE impairs performance in spatial discrimination tasks (Brady et al., 2012; Sanchez et al., 2019) with greater deficits observed when the discrimination involves subtle differences in spatial context (Brady et al., 2012). Importantly, spatial flexibility and discrimination are thought to be reliant on functionally intact hippocampi and have been associated with HPC place cell activity (Behrens et al., 2018; Rubin et al., 2014).

Moderate PAE can also lead to reductions in long-term potentiation (LTP), which is considered to be a mechanism by which increases in synaptic strength support associative learning (Bliss and Lømo, 1973; McNaughton and Morris, 1987). The most prominent reductions in LTP are observed in the dentate gyrus following perforant path stimulation (Brady et al., 2013; Sutherland et al., 1997; Varaschin et al., 2010) and may be related to alterations in NMDA subunit composition (Brady et al., 2013). Importantly, reductions in LTP or NMDA receptor function has been linked to a loss in the spatial stability of HPC place fields (Agnihotri et al., 2004; Barnes, 1979; Barnes and McNaughton, 1980; Barnes et al., 2000; Dieguez and Barea-Rodriguez, 2004; Rotenberg et al., 1996, 2000), and their failure to remap or discriminate spatial contexts (McHugh et al., 2007; discussed in Yassa and Stark, 2011). Thus, it is expected that HPC-dependent spatial discrimination, such as how place cells discriminate directions of locomotion on linear track environments (Battaglia et al., 2004; Frank et al., 2004; McNaughton et al., 1983) or between spatial context changes (Yassa and Stark, 2011), will be disrupted following PAE.

In the present study, we first hypothesized that because PAE impairs spatial behavior and hippocampal function, place cells will have altered spatial tuning, temporal spatial stability, theta rhythmicity, and phase coding in PAE offspring. Secondly, because previous studies report decreased LTP, we hypothesize that place cells from rats with PAE will have decreased directional discrimination on a linear track environment and differential rate and spatial tuning effects following context changes between sessions. We therefore recorded from neurons from HPC CA1 and CA3 of adult rats with moderate PAE (BAC: 60.8 + 5.8 mg/dl (Davies et al., 2019)) that occurred throughout gestation, and saccharin control rats over three environmental conditions: a linear track, an open field cylinder, and an open field cylinder with a local cue rotation (Fig. 1A,B). Our results show that HPC CA3 neurons from PAE rats have reduced spatial tuning and are less likely to display theta phase precession. Secondly, CA1 and CA3 neurons from PAE rats have reduced theta frequencies and are less likely to orthogonalize their firing between directions of travel on the linear track and in context changes between the cylinder sessions compared to control animals.

**Figure 1.**
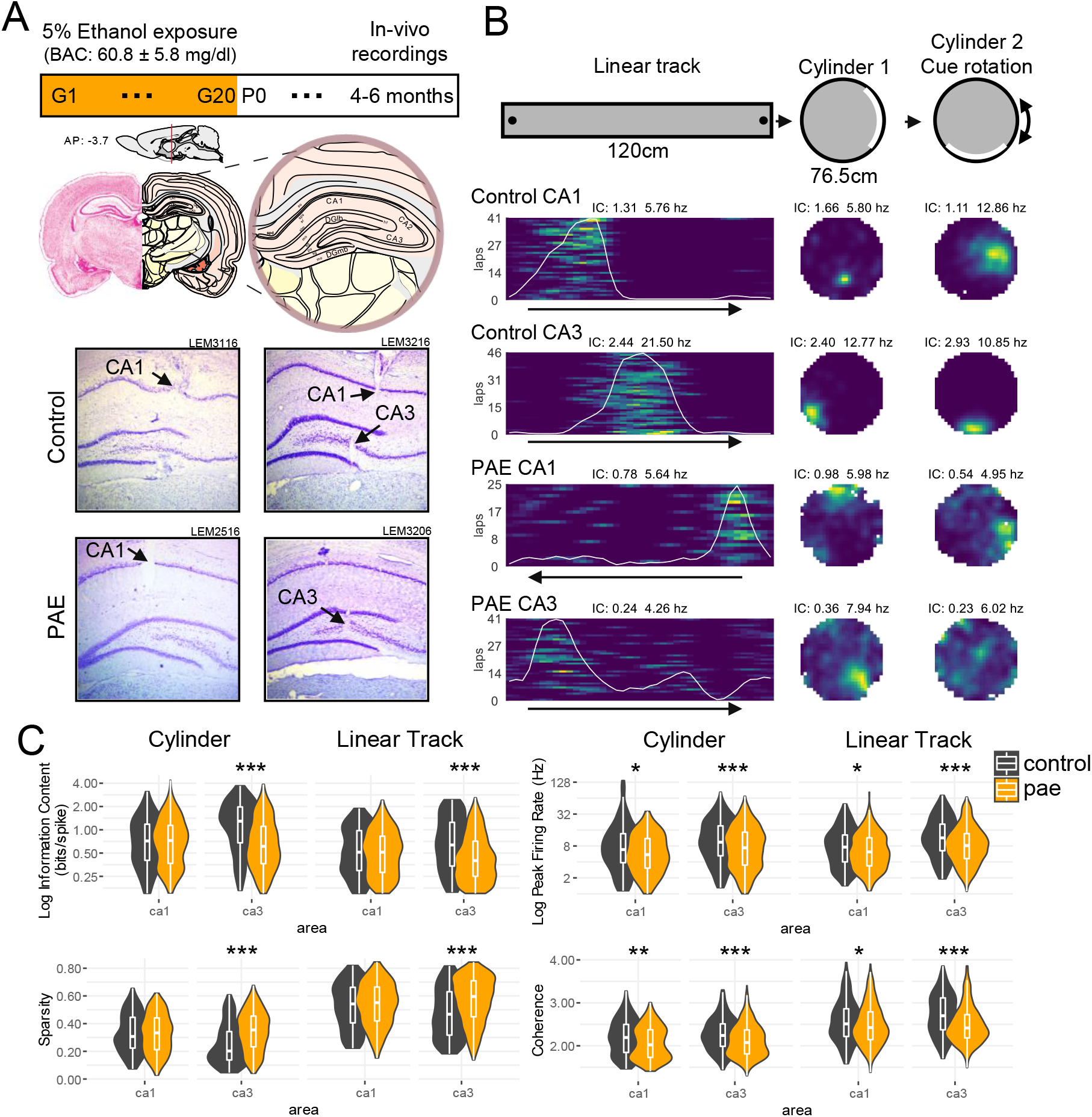
Experimental Protocol, hippocampal place cell examples, & group comparisons. (A) Ethanol exposure paradigm. Top: Ethanol exposure and experimental timeline. Middle: Diagram showing hippocampus in the rat brain (Swanson, 2004). Bottom: Representative Nissl stained histological images (4x magnification) showing electrode penetration (black arrow) in hippocampal CA1 & CA3 from control and PAE rats. (B) Representative place cell examples. Top: Environments for testing spatial tuning of hippocampal place cells. Session 1: 120cm linear track (black dots indicate reward sites), Session 2: 76.5cm open cylinder with salient cue (white), Session 3: 67cm open cylinder with rotated salient cue. Bottom: 4 representative place cells from control and PAE rats in each environment. Information content (IC) and peak firing rate (Hz) are shown above each rate map. Color is scaled with maximum firing rate for each cell and environment. Running direction on the linear track is shown with a black arrow. (C) Violin plots with appended box plot for firing rate, spatial coherence, & spatial tuning measures. Spatial tuning was assessed with spatial information content and sparsity. P-value < 0.05 (*); < 0.01 (**); <0.001 (***).

## Results

### Reduction in spatial tuning of CA3 place cells after PAE

To explore how HPC spatial encoding is affected following moderate PAE, we recorded populations of HPC neurons from 9 control and 8 PAE adult male rats as they ran laps on a linear track or randomly foraged for scattered food in a cylindrical enclosure (Fig. 1A,B). PAE did not disrupt the ability to perform laps on the linear track with no group differences observed in the number of laps completed per session (*p* > 0.05, Wilcoxon rank sum test). There were no significant group differences in locomotor velocity on the linear track (*p* > 0.05, Wilcoxon rank sum test) or in the cylinder (*p* > 0.05, Wilcoxon rank sum test). These findings are consistent with previous studies that have reported unaffected locomotor behaviors following moderate PAE (Brady et al., 2012; Patten et al., 2016).

Despite similar behavioral metrics, impairments in the spatial tuning, measured by spatial information content and sparsity, were evident in PAE place fields. CA3 place fields in PAE animals had significantly lower spatial information and higher sparsity in both linear track and cylinder tests (all *p* < 0.001, Wilcoxon rank sum test, effect sizes (r) ≥ 0.19, Fig. 1C). In contrast, we did not observe group differences in spatial tuning by CA1 place fields.

We also investigated group differences in peak firing rate and spatial coherence which measures the consistency of spiking across the place field. In PAE rats, CA1 and CA3 place cells had significantly diminished peak firing rates in both tasks (all *p* < 0.05, Wilcoxon rank sum test, effect sizes (r) ≥ 0.10, Fig. 1C). Furthermore, CA1 and CA3 firing fields in PAE rats had significantly lower spatial coherence in both tasks (all *p* < 0.05, Wilcoxon rank sum test, effect sizes (r) ≥ 0.08, Fig. 1C). However, it is important to note that the extent of these two CA1 differences were small (effect sizes (r) 0.10 to 0.14). Together, these results indicate that while peak firing and coherence in both CA1 and CA3 were affected by PAE, measures of spatial tuning were affected to a larger extent in CA3.

### Reduced within-session stability by CA1 and CA3 place cells after PAE

To investigate whether spatial tuning was stable over time, we evaluated the within-session spatial stability of place fields on the linear track and cylinder environments (Fig. 2). Within-session stability was measured by calculating the spatial correlation between rate-maps for the first half of the recording session versus the second half. CA1 and CA3 PAE place cells had lower within-session stability compared to control CA1 and CA3 place cells in both linear track and cylinder environments (all *p* < .001, Wilcoxon rank sum test, effect sizes (r) ≥ 0.11, Fig. 2). The decreased spatial stability suggests that PAE negatively affects the ability to create stable spatial representations over relatively short time scales around 20-30 minutes in length.

**Figure 2.**
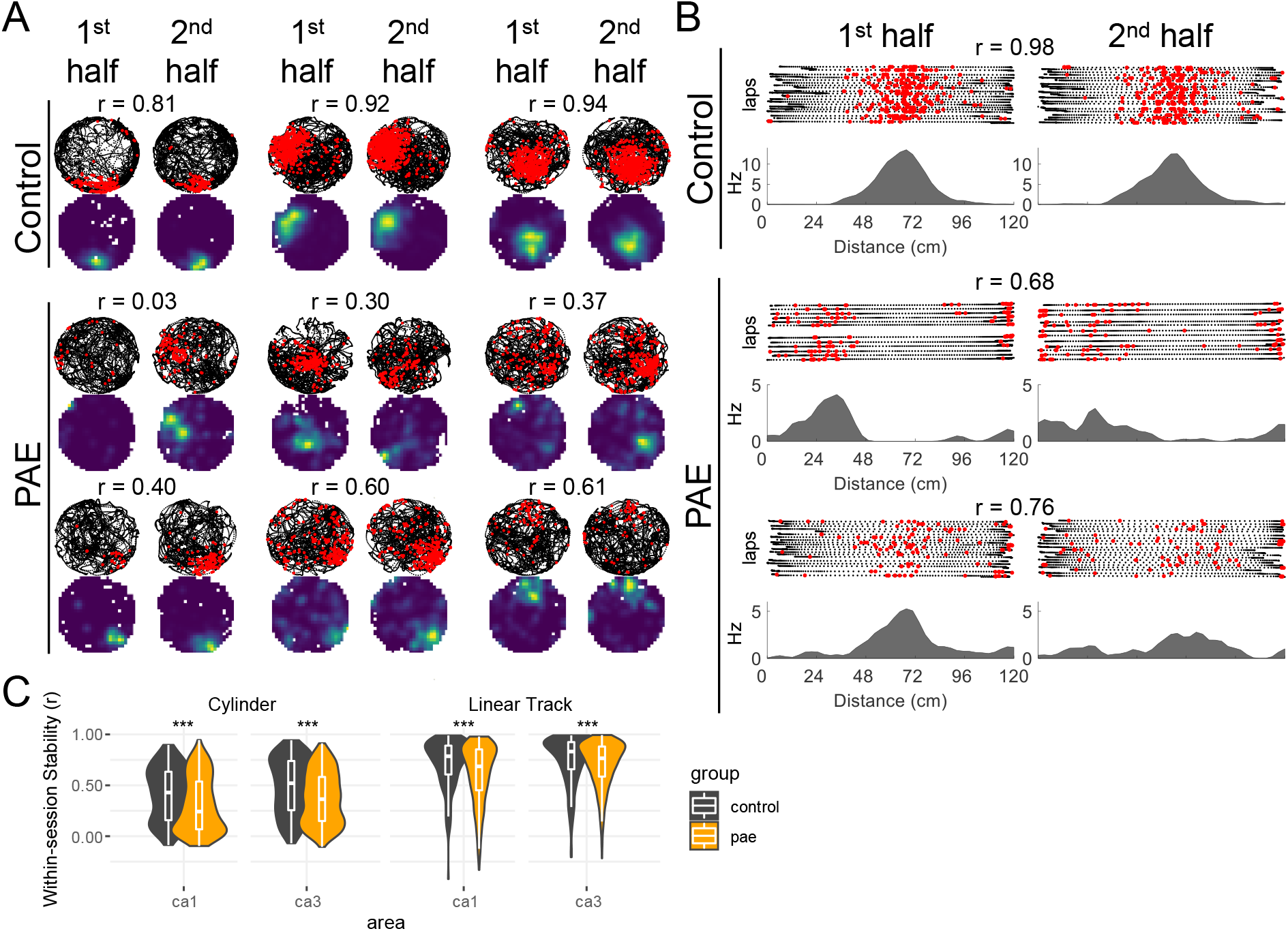
Decreased within-session spatial stability following PAE. (A) Control and PAE example place cells from the first and second halves of their respective recording sessions in the cylinder. Note the decreased stability in the PAE group. Rat’s path (black dots), neuron’s action potentials (red dots). Warm colors represent high firing locations. Within-session spatial correlations are shown above each cell. (B) Control and PAE example place cells from the first and second halves of their respective recording sessions in the linear track. (C) CA1 and CA3 place cells from PAE rats have reduced spatial correlations between the first and second halves of a recording session in both cylinder and linear track environments. P-value < 0.05 (*); < 0.01 (**); <0.001 (***).

### Reduced CA1 and CA3 place cell directionality after PAE

When rodents engage in random foraging in an open-field environment, place cells exhibit location-specific firing that is often independent of the direction of travel through the place field (Muller and Kubie, 1987). However, when the animal’s path is restricted, such as in a linear track, firing rates and peak firing locations significantly differ in opposite directions of travel (Battaglia et al., 2004; McNaughton et al., 1983; Muller et al., 1994). As a result, place cells express directionality in their firing such that their representation of track in the two directions are distinct or orthogonal (Fig. 3A,B). Thus, to determine whether directional firing on the linear track was impaired after PAE, we quantified place cell directionality in two ways: first by correlating rate maps for place fields in each direction of travel and by computing a directionality index which quantifies the distinction in firing rates in each direction. We found that spatial correlations between the two running directions were significantly higher for CA1 PAE place cells (*p* < 0.01, Wilcoxon rank sum test, effect size (r) = 0.17, Fig. 3A-C), but were similar across the two groups for CA3 place cells (*p* > 0.05). This finding indicates that, following PAE, CA1 place cells on the linear track are less likely to spatially orthogonalize. Similarly, the directionality index between the two running directions were significantly lower for CA3 PAE place cells (*p* < 0.001, Wilcoxon rank sum test, effect size (r) = 0.13, Fig. 3A-D) indicating that CA3 place cells are less likely to orthogonalize with respect to their firing rates on the linear track in PAE rats. Together, mechanisms underlying place cell directionality are disrupted suggesting that rats with PAE are unable to form unique complex representations which aid in the disambiguation of different environmental directions.

**Figure 3.**
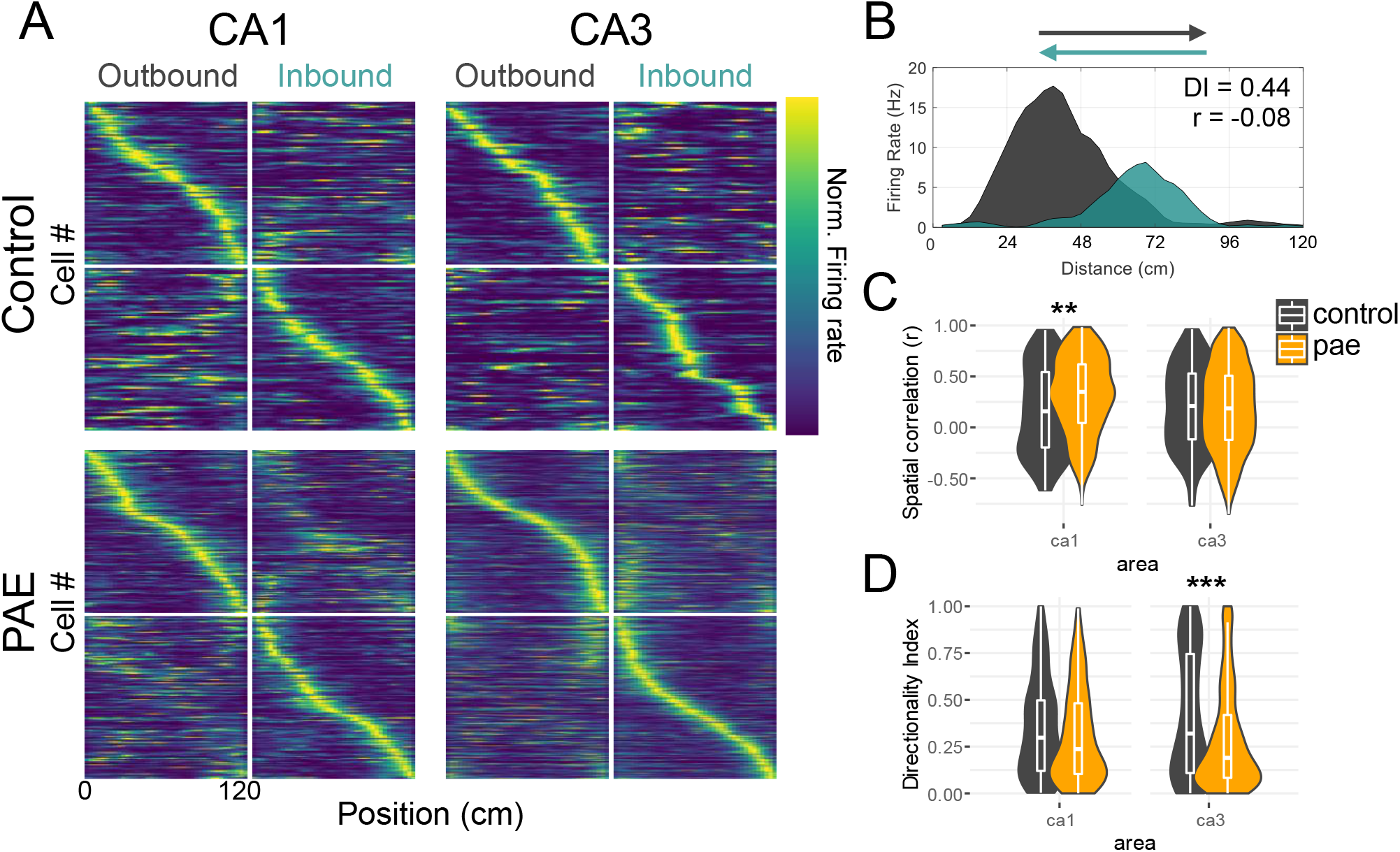
PAE affects place cell directional coding. (A). Place cells sorted by their location of peak firing rate. Each row indicates the firing rate of a single cell as a function of position. Color indicates firing rate normalized by the peak rate across both running for each cell (color bar far right). (B) Example place cell on the linear track showing bidirectional coding noted by the difference in firing rate (directionality index = 0.44) and firing position (spatial correlation = −0.08) between the two running directions. (C) Spatial correlations between rate maps created from the two running directions. Note that PAE CA1 cells are more spatially correlated between the running directions. (D) Directionality index, a metric of firing rate difference, between the running directions. Note that PAE CA3 cells have decreased directionality index values indicating that PAE CA3 cells have similar firing rates between the two running directions. P-value < 0.05 (*); < 0.01 (**); <0.001 (***).

### Dominant control by a proximal cue over CA1 and CA3 place fields after PAE

Given the findings on the linear track, we next sought to determine the sensitivity of PAE place cells to a context change by rotating a cue located along the wall of the open-field cylinder environment (Fig. 4A). We made no attempt to disorient rats between sessions as to not disrupt their internal path integrator thus decreasing the perceived stability of the cue (Knierim et al., 1995; Samsonovich and McNaughton, 1997). A rotational correlation analysis was used to assess the response of place cells following the 90 degree cue rotation by finding the angle at which the two rate maps showed the highest correlation (i.e. rotation angle) (Calton et al., 2003; Harvey et al., 2018; Muller and Kubie, 1987) (Fig. 4A). Cells with correlations below 0.4 were not used in this analysis. The average rotation angle of CA1 place cells was significantly different between PAE and control groups (*F*(1,216) = 6.46, *p* = 0.01, Fig. 4B), though both groups exhibited angular distributions that significantly clustered around the expected shift angle of 90 degrees (Vs ≥ 59.6, *p*s ≤ 0.02). Specifically, CA1 control place cells showed an average rotation angle consistent with cue-averaging whereby an intermediate angle between the first and second cue positions was observed (*θ* = 44.8 degrees, *Z* = 4.00, *p* = 0.01), while CA1 PAE place cells shifted their field locations more closely with the cue (*θ* = 82.84 degrees, *Z* = 24.38, *p* < .001). Similarly, the rotation angle of CA3 place cells was also significantly different between groups (*F*(1,505) = 82.08, *p* < 0.001). CA3 control place cells had a wider distribution of rotation angles that were not clustered around 90 degrees (*θ* = 246.4 degrees, *Z* = 3.01, *p* = 0.048; V = −16.5, p = .98). In contrast, CA3 PAE place cells rotated their fields in correspondence with the cue-shift (*θ* = 73.2 degrees, *Z* = 31.7, *p* < 0.001; V = 107.52, p < .001). Overall, both CA1 and CA3 PAE place cells showed strong dominance of a proximal cue in determining their spatial position within the open field.

**Figure 4.**
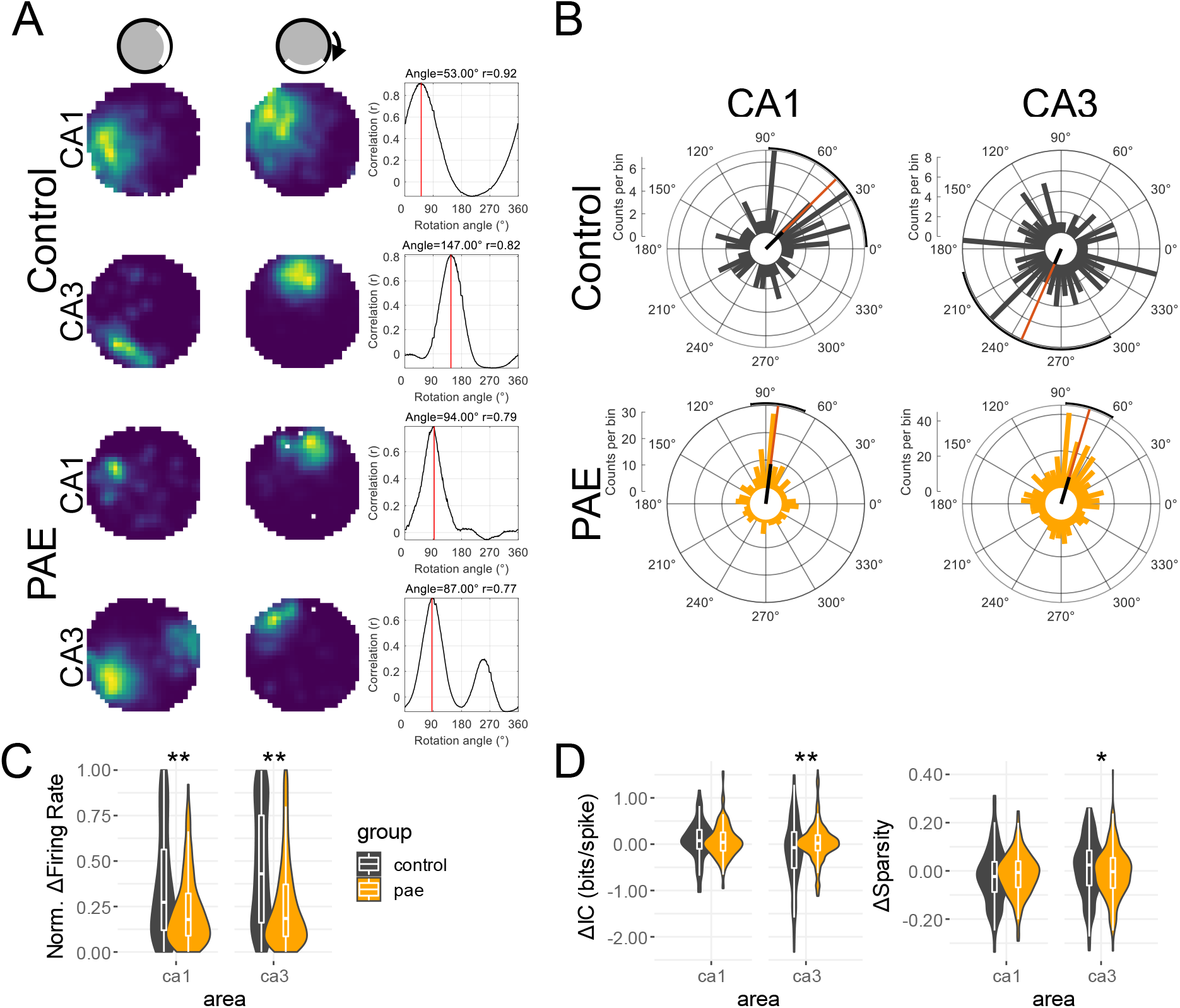
Spatial context orthogonalization. (A) Example place cells from standard cylinder session 1 (left) and the cue rotation session (middle). The right panel shows the rotational correlation analysis where the rate map from the rotation session was rotated in 6 degree bins and a spatial correlation was taken for each correlation. The rotation that resulted in the highest correlation (red line) was assumed to be the directional displacement between the two sessions. (B) Polar histograms showing the distribution of place field rotation following the cue rotation. Note that while control CA1 an CA3 cells have mostly wide distributions, PAE CA1 and CA3 cells display tight distributions clustered around 90 degrees. The orange bar represents mean direction and black bar represents relative mean vector length of each distribution. Polar plots were created using the ‘CircHist’ matlab package (Frederick Zittrell, 2019). (C) Normalized firing rate difference between cylinder session one and two. Control CA1 and CA3 place cells are more likely to exhibit rate remapping compared with PAE cells. (D) Change in spatial tuning (information content and sparsity) between cylinder session one and two. Spatial information and sparsity differences scores were created by subtracting values obtained in session 1 from values in session 2. Note the significant group differences in CA3 cell between both spatial tuning metrics (Δ Information content: control median = −0.07, PAE median = 0.01; Δ Sparsity: control median = 0.02, PAE median = −0.00. P-value < 0.05 (*); < 0.01 (**); <0.001 (***).

We next measured the sensitivity of place cell activity in cue manipulation tests by comparing firing rates and spatial tuning across sessions (Fig. 4C,D). Control cells from CA1 and CA3 demonstrated greater rate remapping between the two sessions compared with PAE cells (CA1: *p* < 0.001, Wilcoxon rank sum test, effect size (r) = 0.20; CA3: *p* < 0.001, Wilcoxon rank sum test, effect size (r) = 0.27, Fig. 4C). Finally, we used difference scores to determine if changes in spatial tuning between the first and cue-shift cylinder sessions would be similar between control and PAE place cells. Spatial tuning of CA3 place cells between the two sessions was differentially changed compared to control cells (Δ spatial information content: *p* < 0.01, Wilcoxon rank sum test, effect size (r) = 0.12; Δ sparsity: *p* < 0.05, Wilcoxon rank sum test, effect size (r) = 0.09, Fig. 4D). Specifically, control cell displayed increased variability between the two conditions. Together, the observations suggest that PAE place cells were less likely to exhibit a change in spatial firing, or more rigidly represented their environment, despite a change in the spatial context.

### Place cell theta modulation and phase precession is disrupted after PAE

Place cells are known to rhythmically fire at theta frequency (4-12Hz) and demonstrate a phenomenon known as “phase precession” when an animal passes through its place field (O’Keefe and Recce, 1993). During phase precession, place cells fires at a progressively earlier phase of the local theta rhythms. Therefore, place cells can simultaneously exhibit rate (place) and phase coding which in effect increases the spatial information contained within any given firing sequence (Huxter et al., 2003). Thus, we next investigated whether spatial impairments after moderate PAE is related to differences in theta-rhythmicity and phase precession of CA1 or CA3 place cells.

We first investigated spike train theta frequencies using a maximum likelihood estimation approach (Climer et al., 2015). We found that the proportion of significant theta rhythmicity was reduced in PAE CA1 and CA3 place cells in the cylinder session (CA1: *X* ^2^ (1, N = 1,691) = 13.92, *p* < 0.001, effect size (V) = 0.02; CA3: *X* ^2^ (1, N = 1,1696) = 32.01, *p* < 0.001, effect size (V) = 0.02) and in PAE CA3 place cells in the linear track (*X* ^2^ (1, N = 1,1646) = 15.55, *p* < 0.001, effect size (V) = 0.01), but not in PAE CA1 cells on the linear track (*p* > 0.05, Fig. 5A) suggesting that environmental and task features in the linear track, such as the two fixed goal sites, may be associated with greater recruitment of theta-rhythmic neurons in CA1. Additionally, PAE CA1 and CA3 had significantly slower peak theta frequencies in both linear track and cylinder environments (all *p* < .001, Wilcoxon rank sum test, effect sizes (r) ≥ 0.16, Fig. 5B-D).

**Figure 5.**
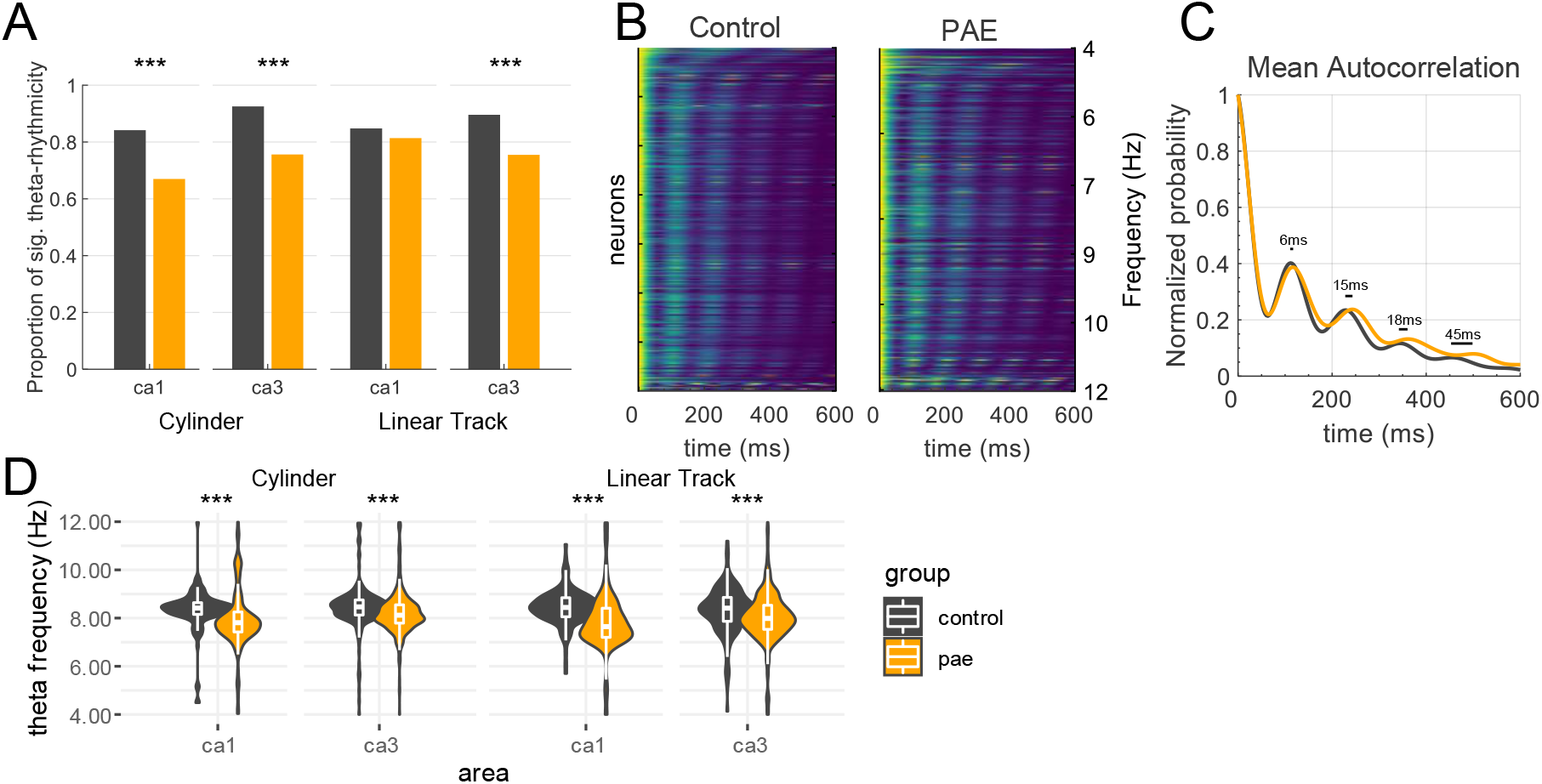
Theta frequency is decreased following PAE. (A) CA1 and CA3 place cells from PAE rats have decreased proportions of significant theta-rhythmicity. (B) Normalized maximum likelihood estimate (MLE) spike-time autocorrelations for control and PAE place cells from CA1 and CA3 and both environments, ordered from bottom to top according to their theta frequency. Warm and cool colors were assigned to the highest and lowest values, respectively, of each autocorrelation. (C) Mean MLE spike-time correlation for all place cells. Note that neurons from the PAE group are approximately 6ms slower on average for each theta cycle. (D) PAE rats have significantly slower spike-time theta frequency across CA1 and CA3 in both linear track and cylinder environments. P-value < 0.05 (*); < 0.01 (**); <0.001 (***).

We next investigated whether PAE place cells exhibited similar phase precession (Fig. 6 A-D). While the strength of theta phase precession was similar between groups in both environment, measured by a circular-linear correlation (all p > 0.09, Fig. 6E), the proportion of classified phase precession was significantly reduced in CA3 from PAE rats on the linear track (*X* ^2^ (1, N = 1,269) = 4.32, *p* = 0.03, effect size (V) = 0.05) and in the cylinder (*X* ^2^ (1, N = 1,301) = 14.44, *p* < 0.001, effect size (V) = 0.01, Fig. 6F). This latter finding suggests that the specific mechanisms underlying CA3 phase precession may be altered following PAE, while mechanisms underlying CA1 phase precession are spared.

**Figure 6.**
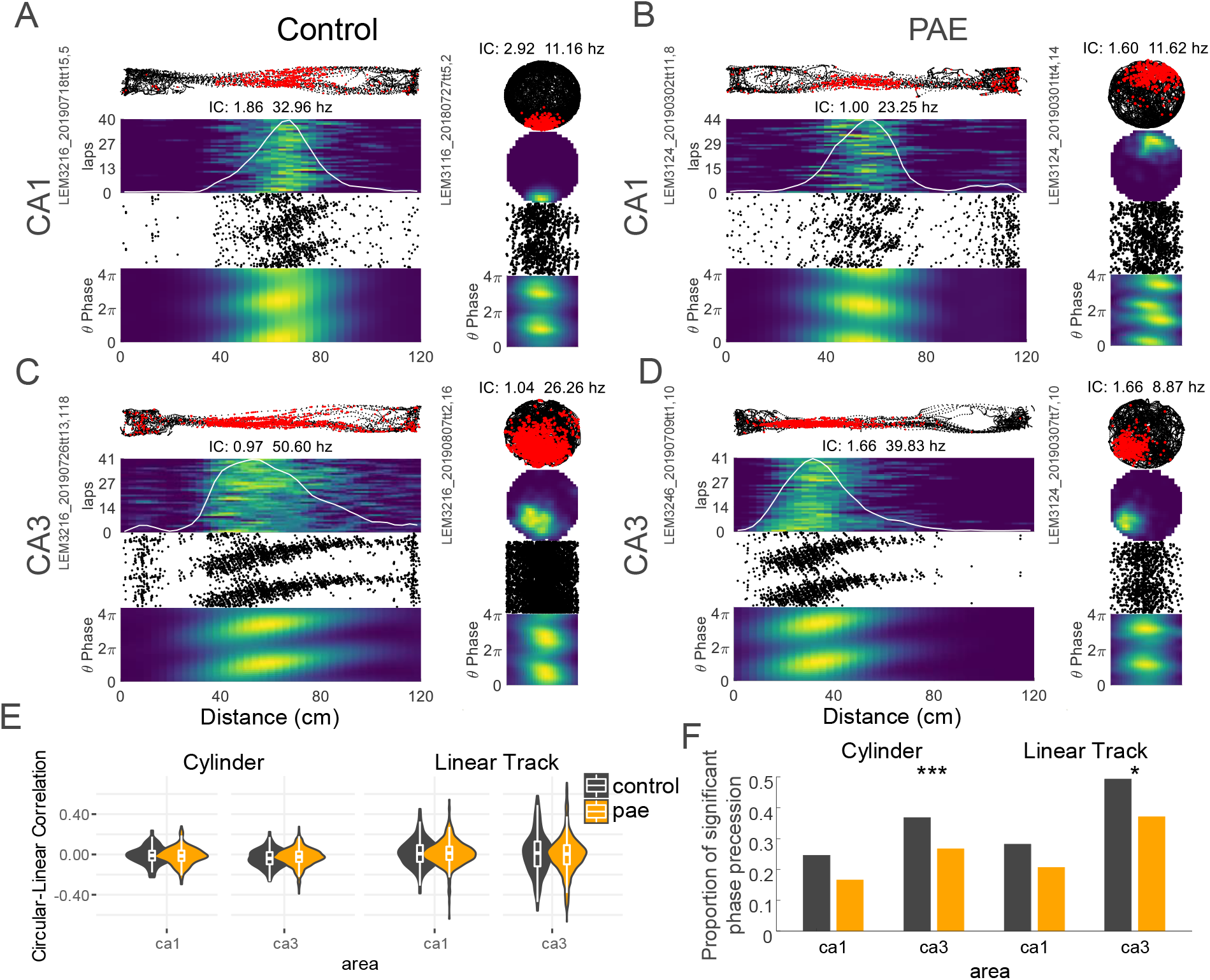
CA3 theta phase precession is affected by PAE. (A-D) Example place cells that demonstrate significant theta-phase precession from control and PAE in both environments. (A-D) Row 1: Rat’s path (black dots), neuron’s action potentials (red dots). Row 2: Rate maps for each lap. Warm colors represent high firing locations. White line represents mean firing rate across laps. Information content (IC) & peak firing rate are listed at the top of each linear track map. Row 3: Distance by theta phase scatter plots. Row 4: Distance by theta-phase rate maps. (E) Circular-linear correlations. Control and PAE place cells had similar circular-linear correlations in both brain regions and in both environments. (F) Proportion of significant theta phase precession. PAE CA3 place cells had significantly reduced proportions of phase precession in both environments. P-value < 0.05 (*); < 0.01 (**); <0.001 (***).

## Discussion

The present study established six specific findings regarding the impact of moderate PAE on hippocampal place cell firing and theta rhythmicity. First, while moderate PAE reduced the consistency of spiking and peak firing rates of place fields across both CA1 and CA3 subregions, the spatial tuning of place fields was disrupted only in CA3. Second, we found that moderate PAE impaired the within-session stability of hippocampal place fields in both CA1 and CA3 regions. Third, moderate PAE disrupted the directionality of CA1 and CA3 place cells on the linear track. Fourth, CA1 and CA3 place fields in PAE animals showed greater control by a proximal landmark. Fifth, we found that PAE slowed peak theta frequencies among CA1 and CA3 place cells and reduced the number of place cell expressing significant theta modulation in the cylinder environment. Finally, PAE reduced the number of place cells exhibiting significant theta-phase precession in CA3.

Several results reported in the present study are likely associated with the observed deficits in hippocampal excitatory signaling at dentate gyrus synapses following PAE. For instance, moderate PAE has been consistently linked to impairments in synaptic plasticity at perforant pathway-dentate gyrus synapses even well into adulthood (Brady et al., 2013; Fontaine et al., 2016; Sutherland et al., 1997; Varaschin et al., 2010). Moreover, a previous study has shown that NMDA subunit composition can be altered in the dentate gyrus of adult mice after moderate PAE (Brady et al., 2013). It has been established that the integrity of synaptic plasticity and NMDA receptor function at perforant pathway synapses is critical for remapping by CA3 place cells (McHugh et al., 2007). Previous studies have also reported that moderate PAE can reduce mGluR5 receptor function in DG (Galindo et al., 2004) and lead to elevated histamine H3 receptor-mediated inhibition of glutamate release from perforant path nerve terminals (Varaschin et al., 2018). Likewise, the integrity of input from the entorhinal cortex, which forms the most prominent source of perforant path axons, is critical for place cell activity (Latuske et al., 2018), and damage to entorhinal cortical cells can produce a similar loss of place field stability and reduction in spatial tuning (Hales et al., 2014; Schlesiger et al., 2018; Van Cauter et al., 2008).

We also found that hippocampal place cells recorded in the PAE group were preferentially controlled by a landmark located along the wall of the cylinder environment. Specifically, when the landmark was rotated by 90 degrees, a large majority of place cells recorded in the PAE group rotated their firing fields in the same angular direction and distance as the cue. In control subjects, place cells were less likley to rotate their firing fields with the landmark but instead demonstated cue-averaging or remapped to largely random locations. The findings supports the conclusion that hippocampal circuitry in PAE animals can establish strong associations with proximal cues, but sensitivity to other contextual stimuli appears more limited. For instance, between cylinder tests (standard and rotation), animals were carefully transported between environments with full access to available distal cues. Thus, stimuli marking the global frame of reference and self-motion cues (e.g., vestibular, motor, etc) remained intact between cylinder sessions. Hippocampal place fields are also influenced by distal and self-motion based cues (Harvey et al., 2018; Jeffery and O’Keefe, 1999; Terrazas et al., 2005; Yoganarasimha et al., 2006) and the conflict between these and the proximal cue rotation may have induced place field remapping in the control group. Indeed, some evidence of some proximal cue influence in the form of cue-averaging was observed for fields in CA1 of the control group (see Fig. 4B). Thus, a weakened capacity to establish associations with distal and/or self-motion cues may be a consequence of moderate PAE. Previous research suggests that information regarding the distal framework and self-motion cues is largely derived from the medial entorhinal cortex input to the hippocampus and indirect sources from other parahippocampal regions (McNaughton et al., 2006; Yoder et al., 2011). As already noted above, perforant path input to the dentate gyrus is weakened in moderate PAE. However, whether deficits are preferential to the medial perforant pathway has not been systematically investigated after moderate PAE. A final point is that an enhancement in proximal cue control may also explain the observed reduction in place field directionality in the PAE group. (Battaglia et al., 2004) reported a similar finding when proximal cues were placed along the surface of a linear track thereby enhancing the salience of the local frame of reference.

While moderate PAE produced several deficits in place cell firing across both CA1 and CA3, the spatial tuning of CA3 place cells were disrupted to a greater extent. In other words, moderate PAE produced noisier CA3 place fields with decreased spatial information relative to control fields in CA3, while CA1 fields from both groups had similar spatial information. In addition, fewer CA3 fields exhibited theta-phase precession after moderate PAE. However, PAE place cells that demonstrated phase precession did so at similar strengths relative to control cells. Although these observations suggest that CA3 might be particularly vulnerable to the effects of moderate PAE, it is important to point out that CA1 and CA3-dependent behaviors are both disrupted after moderate PAE (Patten et al., 2016; Sanchez et al., 2019). In addition, we also found differential effects of moderate PAE on CA1 and CA3 directionality with less discrimination in the spatial position of CA1 place fields and less discrimination in the firing rates of CA3 place cells. As noted earlier, firing rate discrimination by CA3 place cells is dependent on synaptic plasticity in dentate gyrus (McHugh et al., 2007), which is weakened after moderate PAE (Savage et al., 2010). Deficits in CA1 spatial discrimination may also reflect disrupted perforant path input. CA1 place cells also receive direct input from layer III entorhinal cells, but this input is not known to have a strong influence on place cell remapping (Schlesiger et al., 2018) and damage to this projection produces reductions in spatial tuning of CA1 place cells (Brun et al., 2008) which was not observed in the present study.

We established that along with changes in place cell rate coding, we identified changes in place cell theta rhythmicity after moderate PAE. We first found a smaller proportion of PAE CA1 and CA3 place cells modulated by the theta frequency. Secondly, those that were theta-rhythmic had reduced theta frequency compared with control cells. Likewise, this decrease was paralleled with a subtle decrease of firing rates of place cells from PAE rats. This alteration in theta frequency may partially explain previously reported spatial deficits in rodents with PAE (reviewed in Harvey et al., 2019) as the integrity of HPC theta oscillations has been linked to accurate spatial memory (O’Keefe and Nadel, 1978; Winson, 1978). In addition, recent evidence suggests that moderate PAE reduces the number of fast-spiking parvalbumin^+^ GABAergic interneurons throughout the HPC (Madden et al., 2019). Importantly, these interneurons play a vital role in regulating theta-phase precession and HPC theta frequency (Amilhon et al., 2015; Hu et al., 2014; Royer et al., 2012). Consequently, the reduction of theta frequency may be associated with microcircuit changes as a result of the reduced number of GABAergic interneurons. Future studies are needed to determine whether HPC microcircuit function is substantially altered by PAE which would result in theta frequency reductions.

To summarize, the present study establishes a clear linkage between moderate PAE and deficits in spatial and rhythmic firing by hippocampal cell populations. We are unaware of previous work investigating the developmental impact of alcohol on hippocampal place cell activity. In addition, although structural changes to the hippocampus and deficits in hippocampal-dependent behavior have been firmly established after moderate PAE, changes at the level of neural population activity *in vivo* has not received similar systematic investigation. Thus, the findings of the present study represent a critical step in developing a complete multi-level understanding of the neurobiological basis of spatial learning and memory impairments after moderate PAE.

## Methods

### Experimental Model and Subject Details

#### Subjects

Subjects were 17 male Long-Evans rats obtained from the University of New Mexico Health Sciences Center Animal Resource Facility (see breeding protocol below). After weaning, all animals were pair-housed in standard plastic cages with water and food available ad libitum. All cage-mate pairs were matched for age and weight and animals were at least 4 months of age prior to testing. Saccharin-exposed control (n = 9) and prenatal-alcohol-exposed (PAE) (n = 8) rats began linear track and cylinder behavioral training at 4–6 months of age. At this time, rat’s weights were slowly brought down to approximately 90 percent of their ad libitum weight. Lights were maintained on a reverse 12 h:12 h light:dark cycle with lights on at 0900 h. All procedures were approved by the Institutional Animal Care and Use Committee of either the main campus or Health Sciences Center at the University of New Mexico.

#### Breeding and voluntary ethanol consumption during gestation

Breeding procedures were conducted at the University of New Mexico Health Sciences Animal Resource Facility. Three to four-month-old female breeders (Harlan Industries, Indianapolis, IN) were single housed in standard plastic cages and placed on a 12-hour reverse light: dark cycle (lights on from 2100-0900 h) and kept at 22 °C with ad libitum food and water. Following a one-week acclimation period in the animal facility, the breeders were exposed to a voluntary ethanol drinking paradigm (Fig. 1A). Female rats were provided 0.066% (w/v) saccharin in tap water from 10:00 to 14:00 h (4 h) each day. On Days 1–2, the saccharin water contained 0% ethanol, on days 3–4 saccharin water contained 2.5% ethanol (v/v), and on day 5 and thereafter, saccharin water contained 5% ethanol (v/v). The daily four-hour consumption of ethanol was monitored for at least two weeks and the mean daily ethanol consumption was determined for each female. After two weeks of daily ethanol consumption, females that drank at levels less than one standard deviation below that of the entire group mean were removed from the study (∼12–15% of all female rats). The remaining females were then assigned to either a saccharin control or 5% ethanol drinking group. These breeding females were matched such that the mean pre-pregnancy ethanol consumption by each group was similar. As a result, dams of both groups experience equivalent preconceptual exposure to ethanol. Lastly, the female breeders were nulliparous and were not used in multiple rounds of breeding, while the male rats were experienced breeders.

Female rats were matched with a male breeder rat until pregnancy was verified, based on the presence of a vaginal plug. There was no ethanol consumption during breeding. Beginning on gestational day 1, the rat dams were given access to saccharin (Sigma Life Sciences, St. Louis, Missouri) water containing either 0% (v/v) or 5% (v/v) ethanol (Koptec, King of Prussia, Pennsylvania) for four hours a day, from 10:00 to 14:00 h. The volume of the 0% ethanol saccharin water provided to the control group was matched to the mean volume of the 5% ethanol saccharin water consumed by the ethanol group. During gestation and including the four-hour ethanol/saccharin drinking period, rats were provided with ad libitum water and rat chow (Teklad global soy protein-free extruded food 2920). Daily ethanol consumption was recorded for each rat dam (Table S1). The daily mean ethanol consumption throughout pregnancy was 1.96 ± 0.14 g/kg) and did not vary significantly during each of the three weeks of gestation. In a separate set of rat dams, this level of ethanol consumption has been shown to produce a mean peak maternal serum ethanol concentration of 60.8 ± 5.8 mg/dl (Davies et al., 2019). Daily ethanol consumption ended at birth, and the litters were weighed and culled to 10 pups. As shown in Table S1, this moderate exposure paradigm did not affect maternal weight gain, offspring litter size, or birth weight for the offspring used in these studies. Experimental offspring used in these studies were generated from nine different saccharin control and eight different ethanol-consuming rat dams bred in four separate breeding rounds.

#### Behavioral training

Each animal was exposed to each maze for several days. On the linear track, animals were trained to continuously alternate between the two ends. Stop locations were rewarded with 1/8 piece of a fruit-loop if animals completed a successful lap without stopping or alternating the incorrect direction. This training continued until 40 laps could be completed within 15 minutes. For the cylinder, animals were trained to randomly forage for fruit-loop pieces that were semi-randomly dropped into the environment until all locations were sufficiently sampled.

### Method Details

#### Surgical procedures

Rats were anesthetized with isoflurane and positioned in a stereotaxic apparatus (David Kopf Instruments) with bregma and lambda set at the same D-V plane. The scalp was retracted, and a hole was drilled above the hippocampus. Six additional holes were drilled in the frontal, parietal, and occipital bones to hold jeweler’s screws with a single screw in the parietal plate used for grounding. With the electrode bundle (see below for fabrication details) positioned dorsal to hippocampus (Fig. 1A), the drive assemblies were fastened to the skull and jeweler’s screws with dental acrylic. The scalp was sutured around the electrode drive and the wound was covered with Neosporin. Buprenorphine was administered postoperatively and the animal was allowed to recover at least 1 week before recording.

#### Apparatus

The cylinder (diameter: 76.5 cm, height: 40 cm) was positioned on a black plastic surface, and a white cue card covered 100 degrees of the wall surface (Fig. 1B). The linear track (length: 120 cm, width: 9 cm, height: 10 cm) was positioned relative to polarizing distal cues (white cue cards) as to make each running direction distinct. The entire recording area was located within a large Faraday cage to reduce electrical noise. Eight battery-powered LED lights mounted on the ceiling provided illumination.

#### Recording Procedures

Single units and local field potentials (LFP) were recorded from the CA1 and CA3 regions of the dorsal hippocampus using 8 to 16 tetrodes each of which consisting of four-channel electrodes constructed by twisting together strands of insulated 12-17 um nichrome wire (Harvey et al., 2018).

After surgery, tetrodes were slowly advanced into the CA1 area of the hippocampus. During tetrode advancement and during recordings, the electrode assembly was connected to a multichannel, impedance matching, unity gain preamplifier headstage. The output was routed to a data acquisition system with 64 digitally programmable differential amplifiers (Neuralynx, Tucson, AZ, USA). Spike waveforms above a threshold of 30-40 µV were time-stamped, digitized at 32 kHz, and later imported into MClust for spike sorting. Continuous signal from each channel were collected for sessions spike sorted with kilosort2. The rat’s position was tracked and sampled at 30 Hz by recording the position of light-emitting diodes that were placed above the head.

Rats were screened each day on a pedestal for identifiable units and HPC theta rhythm visible in the local field potentials. Once units and theta rhythm were found a typical recording began with first ~40 laps on a linear track (~10-15 minutes), followed by an open field cylinder session, and finished with final open field cylinder session with the local cue rotated 90 degrees (Fig. 1B). Linear track sessions in which the rat made less than 15 laps were not included in any analysis. In the cylinder sessions, rats randomly foraged for fruit-loops until the environment was fully sampled (~15-30 minutes). Depending on the rat’s motivation, some sessions were limited to either a single linear track or a linear track and one cylinder session. After each daily recording session, tetrodes were advanced approximately 62μm in order to record different neurons each day. Each rat performed 32 daily sessions on average.

#### Histology

Rats were euthanized by transcardial perfusion with normal saline followed by 10% Formalin, and brains were then placed in Formalin for 24 hours to ensure adequate fixation. Brains were then placed in 20% sucrose for cryoprotection before they were sectioned at 40 mm on a freezing microtome. Brain sections were mounted on gelatin-coated or charged microscope slides and stained with Cresyl violet (Fig. 1A). Electrode position at the time of each recording was estimated relative to the site of the final electrode tip location. Electrode tips were marked by applying constant current (4 mA for 4 s) through a wire of each tetrode. Our analyses included only sessions where the electrode tip was estimated to be located within CA1 and CA3 regions. It is important to note the possibility that a small number of recordings may have included granule cells or mossy cells from the dentate subregion, given its proximity to CA3.

### Quantification and Statistical Analysis

#### Data analysis

Data analysis was performed by importing position data, LFP data, and spike data into Matlab and by further processing the data with custom-written software. Statistical analysis was performed in Matlab and R.

#### Spike sorting

Spike sorting was performed first by using unsupervised methods klustakwik (http://klustakwik.sourceforge.net/) or kilosort2 (https://github.com/MouseLand/Kilosort2) with manual refinement conducted in MClust (https://github.com/adredish/MClust-Spike-Sorting-Toolbox) or Phy (https://github.com/cortex-lab/phy). Autocorrelation and cross-correlation functions were used as single unit identification criteria for both methods. Active neurons, defined as having at least 100 spikes and a peak firing rate of 1Hz, were considered for analysis.

#### Spatial rate map construction

First, spikes that occurred when running speeds were less than 3cm/s were excluded. Then occupancy normalize firing rate maps were constructed using 3×3cm spatial bins. Maps were then smoothed with a 5×5 standard deviation Gaussian kernel. For linear track tuning curves, the x coordinate was binned as above while the y coordinate was compressed into a single bin.

#### Spatial information

Spatial information content for each cell, the spatial information content was calculated as:

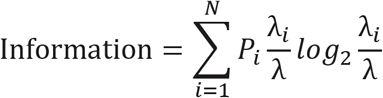

Where the environment is divided into spatial bins *i* = 1, …, *N*, *P*_*i*_ is the occupancy probability of bin *i*, *λ*_*i*_ is the mean firing rate for bin *i*, and *λ* is the overall mean firing rate of the cell (Skaggs and McNaughton, 1996).

#### Sparsity

Sparsity for each cell was calculated as:

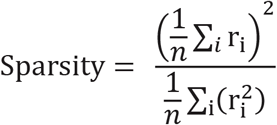

Where *i* is a spatial bin and *r*_*i*_ is the firing rate of the cell in bin *i* of an environment containing a total or n spatial bins (Ahmed and Mehta, 2009). A sparsity value of 1 implies no sparseness. A sparsity value approaching 0 is indicative of maximal sparseness and implies a greater amount of spatial information in each spike emitted by that cell.

#### Spatial correlation

The spatial similarity of place fields across two conditions was calculated using Pearson’s correlation. The correlation coefficient was calculated by comparing the firing rates between all pixels at corresponding locations.

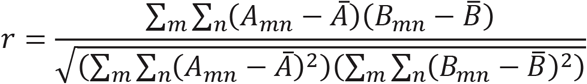

#### Spatial coherence

Spatial coherence was computed as the Fisher z-transformation (the arctangent) of the correlation between the firing rate of each spatial bin and the average firing rate of the neighboring 3-8 spatial bins (Butler et al., 2019).

#### Directionality Index

The directionality of a cell on the linear track was defined as:

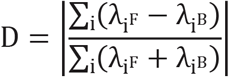

Where 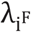 is the firing rate in bin i in the forward (or outbound) running direction and 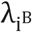 is the firing rate in the backwards (or inbound) running direction (Ravassard et al., 2013). This equation was also used to compare rate differences between the two cylinder sessions.

#### Place cell classification

From neurons recorded during the linear track sessions, the set of heuristics used to identify a ‘place field’ were the following 1) Minimum peak firing rate of 1 Hz, 2) Minimum field width of 9 cm, 3) Maximum field width of 78 cm (65 % of track length), 4) At least 15 laps with consistent behavior, 5) At least 100 spikes, 6) Minimum spatial information content of 0.15 bits/spike. Given these criteria, 24.6% (1459 of 5927) of active (>100 spikes & > 1Hz peak rate) HPC neurons had at least one place field with a mean of 1.26 ± 0.01 per neuron. No limit was set on the number of place fields single neurons could have and metrics were obtained from the field with the highest firing rate. The median peak firing rate was 7.52 ± 0.32 Hz, and the mean field width was 45.93 ± 0.40 cm. The start and stop position for each field were taken as the spatial bins where the firing rate drops below 20% of the peak firing rate.

From neurons recorded in the cylinder sessions, a similar set of heuristics used to identify a ‘place field’ were the following 1) Minimum peak firing rate of 1 Hz, 2) Minimum field width of 9 cm, 3) Maximum field width of 49.7 cm (65 % of cylinder diameter), 4) At least 100 spikes, 5) Minimum spatial information content of 0.15 bits/spike. Given these criteria, 24.1% (1554 of 6448) of HPC active neurons recorded during the first cylinder session had at least one place field with a mean of 1.74 ± 0.02 fields per neuron. While there were no limits set on the number of place fields single neurons could have, place cell metrics were obtained from the field with the highest firing rate. The median peak firing rate was 7.25 ± 0.33 Hz, and the mean field width was 31.71 ± 0.25 cm. The field boundaries were drawn using Matlab’s contourc function following a k-means clustering procedure which separated low and high firing locations.

#### Instantaneous theta phase, firing phase

A 3^rd^ order Butterworth bandpass filter (4–12-Hz) was applied to the raw LFP signal from a single channel on each tetrode selected for phase estimation. The instantaneous theta phase was obtained from the Hilbert transform of the filtered signal. Spike and LFP timestamps were then used to linearly interpolate firing phase from these values.

#### Theta phase precession

One-dimensional phase precession of place cells in the linear track was quantified using a circular-linear correlation between theta firing phase from each place cell and the linear x coordinate as the rat passed through a place field (Jammalamadaka and Sengupta, 2001; Kempter et al., 2012). Phase precession was quantified for multiple place fields from a single cell. However, metrics for comparison were only taken from the place field with the highest peak firing rate.

Two-dimensional phase precession of place cells in the cylinder was computed using ‘pass-index’ (https://github.com/jrclimer/Pass_Index) (Climer et al., 2013). Rate maps were smoothed with a pseudo-Gaussian kernel with a five-bin standard deviation. The value at each bin was then percentile normalized between 0 and 1 which resulted in the field index map. The trajectory of the animal was sampled evenly along the arc-length of the trajectory at as many points as there were position tracking samples (30 Hz). The nearest bins were then found by minimizing the difference between the x and y positions and the center of the bins via the MATLAB function: bsxfun. The omnidirectional pass index was than computed on the field index maps which resulted in a −1 to 1 vector where −1 represents when the animal was at the beginning of the place field, 0 represents the center, and +1 represents the end of the place field. A circular-linear correlation between theta firing phase from each cell and the above vector to quantification of phase precession (Jammalamadaka and Sengupta, 2001; Kempter et al., 2012).

#### Spike-train theta rhythmicity

We assessed the properties of theta-rhythmicity, such as mean frequency of the theta modulation, rhythmicity magnitude, and approximated best fit autocorrelograms, using a maximum likelihood estimation approach (Climer et al., 2015 available at https://github.com/jrclimer/mle_rhythmicity).

#### Statistical Analysis

All statistical tests used were two-sided, and the significance threshold for all tests was set at *p* < 0.05.

For each sample distribution, a Kolmogorov-Smirnov (KS) test was used to test the null hypothesis that the *z*-scored sample was derived from a standard normal distribution. If the KS null hypothesis failed rejection, a one-sample *t* test was used to test the sample mean against zero. Otherwise, a sign test was used to test the sample median against zero.

For each between-group comparison, a two-sample *t* test was used to test equality of means only if both sample distributions failed KS test rejection; otherwise, a Wilcoxon signed-rank test was used to test the equality of medians. Effect size (r) for Wilcoxon signed-rank test was calculated with r = Z/√ n (Cooper and Hedges, 1993).

Between group comparison of proportions were computed using chi-square tests, and effect sizes for these comparisons were calculated with Cramér’s V.

Angular data were assessed using the CircStats2012a matlab toolbox. Verification of von mises distributions were performed using Rayleigh Tests, while a Watson-Williams multi-sample test for equal means was used to compare mean directions between groups.

All tests are shown in Table S2.

## Acknowledgments

The authors thank Dr. Suzy Davies and Dr. Jennifer Wagner for supervising the moderate PAE paradigm and Andre Moezzi, Nicole Graham, Kiana Lujan, Ella Rappaport, and Chloe Puglisi for their assistance with the rodent husbandry procedures. The research reported in this publication was supported by National Institute on Alcohol Abuse and Alcoholism grants R21 AA024983 and P50 AA022534 to BJC, DAH, and DDS, and T32 AA014127-15 to REH.

## Author Contributions

RH, DS, DH, and BC conceptualized experiments. BC, RH, DH, and DS contributed funding. BC contributed lab resources and supervision. DS created the animal model and directed the production of experimental offspring. RH and LB collected and analyzed data with input from BC. RH, LB, and BC wrote the initial draft of the paper with DS and DH providing review and editing.

## Data and software availability

The dataset will be made available upon request to corresponding author. Custom code is available at https://github.com/ryanharvey1/ephys_tools

## Supplemental information

**Supplemental table 1.** Impact of 5% ethanol consumption on maternal weight gain, litter size, and pup birthweight. All values are represented as mean ± standard error of the mean.

|  | Saccharin Control (n = 9) | 5% Ethanol Group (n = 8) |
| --- | --- | --- |
| Daily four-hour 5% ethanol consumption | NA | $1.96 \pm 0.14$ g/kg/d |
| Maternal weight gain during pregnancy | $105 \pm 7$ g | $120 \pm 7$ g |
| Litter size | $10 \pm 0.8$ live births | $11.9 \pm 0.8$ live births |
| Pup birth weight | $7.79 \pm 0.38$ g | $7.84 \pm 0.56$ g |

**Supplemental table 2.** Statistics table showing each test conducted. n = sample sizes, c = control, p = PAE

| Dependent Variable | n | test | Test statistic | P-value | 95 percent CI | effect measure | Effect size |
| --- | --- | --- | --- | --- | --- | --- | --- |
| Number of laps | C: 64<br>P: 111 | Wilcoxon rank sum test | 4182 | 0.05036 | -5.929951e-06 4.000034 | r | 0.1479258 |
| Linear track velocity | C: 64<br>P: 111 | Wilcoxon rank sum test | 3265 | 0.3748 | -1.0228582 0.4127174 | r | 0.06709473 |
| Cylinder velocity | C: 62<br>P: 95 | Wilcoxon rank sum test | 2755 | 0.4962 | -0.9127863 0.4603628 | r | 0.054308 |
| Spatial Information Content linear track CA1 | C: 220<br>P: 421 | Wilcoxon rank sum test | 48178 | 0.4015 | -0.02831160 0.07574237 | r | 0.03313617 |
| Spatial Information Content linear track CA3 | C: 215<br>P: 1049 | Wilcoxon rank sum test | 147395 | 1.234e-12 | 0.1347696 0.2646247 | r | 0.1997467 |
| Spatial Information Content cylinder CA1 | C: 162<br>P: 306 | Wilcoxon rank sum test | 24899 | 0.9356 | -0.08226948 0.09065277 | r | 0.003736075 |
| Spatial Information Content cylinder CA3 | C: 247<br>P: 839 | Wilcoxon rank sum test | 145298 | <2.2e-16 | 0.4031402 0.6303361 | r | 0.2919239 |
| Sparsity linear track CA1 | C: 220<br>P: 421 | Wilcoxon rank sum test | 45640 | 0.7636 | -0.03158489 0.02310201 | r | 0.01187934 |
| Sparsity linear track CA3 | C: 215<br>P: 1049 | Wilcoxon rank sum test | 79562 | 9.765e-12 | -0.12377246 -0.06886454 | r | 0.1915439 |
| Sparsity cylinder CA1 | C: 162<br>P: 306 | Wilcoxon rank sum test | 24907 | 0.931 | -0.02812478 0.02927698 | r | 0.004001751 |
| Sparsity cylinder CA3 | C: 247<br>P: 839 | Wilcoxon rank sum test | 64267 | <2.2e-16 | -0.13053193 -0.08512575 | r | 0.2755911 |
| Peak firing rate linear track CA1 | C: 220<br>P: 421 | Wilcoxon rank sum test | 51947 | 0.01134 | 0.2178625 1.8636384 | r | 0.1000118 |
| Peak firing rate linear track CA3 | C: 215<br>P: 1049 | Wilcoxon rank sum test | 137478 | 4.026e-07 | 1.613853 3.863249 | r | 0.1425403 |
| Peak firing rate cylinder CA1 | C: 162<br>P: 306 | Wilcoxon rank sum test | 28043 | 0.01931 | 0.163791 2.150657 | r | 0.1081469 |
| Peak firing rate cylinder CA3 | C: 247<br>P: 839 | Wilcoxon rank sum test | 120978 | 6.149e-05 | 0.9750111 2.9797256 | r | 0.1215923 |
| Coherence linear track CA1 | C: 220<br>P: 421 | Wilcoxon rank sum test | 50907 | 0.03893 | 0.004486582 0.158407123 | r | 0.08155845 |
| Coherence linear track CA3 | C: 215<br>P: 1049 | Wilcoxon rank sum test | 145828 | 1.2e-11 | 0.1763259 0.3168864 | r | 0.1907075 |
| Coherence cylinder CA1 | C: 162<br>P: 306 | Wilcoxon rank sum test | 29192 | 0.001551 | 0.04691421 0.20635266 | r | 0.1463047 |
| Coherence cylinder CA3 | C: 247<br>P: 839 | Wilcoxon rank sum test | 126441 | 1.38e-07 | 0.0936456 0.2023938 | r | 0.1598539 |
| Stability linear track CA1 | C: 220<br>P: 421 | Wilcoxon rank sum test | 56229 | 8.362e-06 | 0.03999786 0.11008619 | r | 0.1759899 |
| Stability linear track CA3 | C: 215<br>P: 1049 | Wilcoxon rank sum test | 132852 | 3.806e-05 | 0.02572039 0.07431632 | r | 0.1158551 |
| Stability cylinder CA1 | C: 162<br>P: 306 | Wilcoxon rank sum test | 30629 | 2.699e-05 | 0.05554166 0.15883213 | r | 0.1940268 |
| Stability cylinder CA3 | C: 247<br>P: 839 | Wilcoxon rank sum test | 129414 | 2.615e-09 | 0.08134618 0.16137396 | r | 0.1806761 |
| Spatial correlation linear track CA1 | C: 220<br>P: 421 | Wilcoxon rank sum test | 20555 | 0.0001676 | -0.24337755 -0.07667529 | r | 0.1715964 |
| Spatial correlation linear track CA3 | C: 215<br>P: 1049 | Wilcoxon rank sum test | 67057 | 0.8464 | -0.06421564 0.08014783 | r | 0.006195165 |
| Directionality index linear track CA1 | C: 220<br>P: 421 | Wilcoxon rank sum test | 28819 | 0.06507 | -0.002519559 0.085009707 | r | 0.08411345 |
| Directionality index linear track CA3 | C: 215<br>P: 1049 | Wilcoxon rank sum test | 81396 | 1.495e-05 | 0.0450416 0.1406268 | r | 0.13844 |
| Mean angle of Rotation cylinder CA1 | C: 96<br>P: 148 | Watson Williams | 6.4663 | 0.0117 | na | na | na |
| Mean angle of rotation cylinder CA3 | C: 108<br>P: 398 | Watson Williams | 82.0876 | <2.2e-16 | na | na | na |
| Distribution of angles CA1 control | 96 | Rayleigh test of nonuniformity | 4.0091 | 0.0176 | na | na | na |
| Distribution of angles CA3 control | 108 | Rayleigh test of nonuniformity | 24.3806 | 9.4667e-12 | na | na | na |
| Distribution of angles CA1 pae | 148 | Rayleigh test of nonuniformity | 3.0124 | 0.0488 | na | na | na |
| Distribution of angles CA3 pae | 398 | Rayleigh test of nonuniformity | 31.6774 | 9.4583e-15 | na | na | na |
| Distribution of angles around 90 CA1 control | 96 | V test of nonuniformity | 11.7198 | 0.0230 | na | na | na |
| Distribution of angles around 90 CA3 control | 108 | V test of nonuniformity | 59.6025 | 2.1244e-12 | na | na | na |
| Distribution of angles around 90 CA1 pae | 148 | V test of nonuniformity | -16.5311 | 0.9878 | na | na | na |
| Distribution of angles around 90 CA3 pae | 398 | V test of nonuniformity | 107.5290 | 1.2434e-14 | na | na | na |
| Normalized delta firing rate CA1 | C: 96<br>P: 148 | Wilcoxon rank sum test | 29859 | 1.939e-05 | 0.04492751 0.13175392 | r | 0.199393 |
| Normalized delta firing rate CA3 | C: 108<br>P: 398 | Wilcoxon rank sum test | 142264 | <2.2e-16 | 0.1300228 0.2227179 | r | 0.2758055 |
| delta spatial information content CA1 | C: 96<br>P: 148 | Wilcoxon rank sum test | 7512 | 0.2792 | -0.03648802 0.13537247 | r | 0.06857867 |
| Delta spatial information content CA3 | C: 108<br>P: 398 | Wilcoxon rank sum test | 26962 | 0.003245 | -0.24146435 -0.04725801 | r | 0.1205709 |
| Delta sparsity CA1 | C: 96<br>P: 148 | Wilcoxon rank sum test | 6532 | 0.4595 | -0.03638187 0.01478238 | r | 0.0468788 |
| Delta sparsity CA3 | C: 108<br>P: 398 | Wilcoxon rank sum test | 36150 | 0.02879 | 0.002553735 0.044025642 | r | 0.08955616 |
| Proportion of sig. theta cylinder ca1 | C: 162<br>P: 306 | Chi-squared | 13.9286 | 1.8988e-04 | na | V | 0.0202 |
| Proportion of sig. theta cylinder ca3 | C: 247<br>P: 839 | Chi-squared | 32.0124 | 1.5319e-08 | na | V | 0.0189 |
| Proportion of sig. theta linear track ca1 | C: 220<br>P: 421 | Chi-squared | 0.8192 | 0.3654 | na | V | 0.0010 |
| Proportion of sig. theta linear track ca3 | C: 215<br>P: 1049 | Chi-squared | 15.5542 | 8.0171e-05 | na | V | 0.0094 |
| Theta frequency cylinder ca1 | C: 162<br>P: 306 | Wilcoxon rank sum test | 14755 | 7.215e-10 | 0.4199294 0.7042946 | r | 0.3605629 |
| Theta frequency cylinder ca3 | C: 247<br>P: 839 | Wilcoxon rank sum test | 75874 | 4.921e-10 | 0.2070257 0.3894830 | r | 0.2267278 |
| Theta frequency linear track ca1 | C: 220<br>P: 421 | Wilcoxon rank sum test | 21929 | 5.534e-13 | 0.5360641 0.8692292 | r | 0.3785056 |
| Theta frequency linear track ca3 | C: 215<br>P: 1049 | Wilcoxon rank sum test | 51565 | 1.983e-05 | 0.1852935 0.4803838 | r | 0.1589059 |
| Proportion of sig. phase precession cylinder ca1 | C: 162<br>P: 306 | Chi-squared | 2.4898 | 0.1146 | na | V | 0.072 |
| Proportion of sig. phase precession cylinder ca3 | C: 247<br>P: 839 | Chi-squared | 14.183 | 0.0001658 | na | V | 0.096 |
| Proportion of sig. phase precession linear track ca1 | C: 220<br>P: 421 | Chi-squared | 2.49 | 0.1146 | na | V | 0.06 |
| Proportion of sig. phase precession linear track ca3 | C: 215<br>P: 1049 | Chi-squared | 4.7067 | 0.03005 | na | V | 0.06 |
| Circular-linear correlation cylinder ca1 | C: 162<br>P: 306 | Wilcoxon rank sum test | 25599 | 0.5594 | -0.01083565 0.02052675 | r | 0.02698276 |
| Circular-linear correlation cylinder ca3 | C: 247<br>P: 839 | Wilcoxon rank sum test | 95289 | 0.05462 | -0.0241500306 0.0002705691 | r | 0.05832034 |
| Circular-linear correlation linear track ca1 | C: 220<br>P: 421 | Wilcoxon rank sum test | 46199 | 0.9604 | -0.01994809 0.01890835 | r | 0.001960668 |
| Circular-linear correlation linear track ca3 | C: 215<br>P: 1049 | Wilcoxon rank sum test | 116127 | 0.4909 | -0.01713642 0.03590418 | r | 0.01937648 |

